# Pheromone Receptors Help *Cnaphalocrocis medinalis* Avoid Competition in Rice Fields

**DOI:** 10.1101/2021.05.27.445942

**Authors:** Jianjun Cheng, Yongle Zhang, Yongjun Du

## Abstract

*Cnaphalocrocis medinalis* (Guene’e) (Lepidoptera: Pyralidae) is one of the most important insect pests that attack the rice crop, *Oryza sativa* L., in China, feeding on rice leaves. *Chilo suppressalis* and *Sesamia inferens* are two common insects living within the same ecological system that feed on rice stalks. Their behavior could affect *C. medinalis*’s choice of oviposition place, so we tested the electroantennogram (EAG) response of *C. medinalis* to a conspecific sex pheromone (Z11-18:OH; Z11-18:Ald; Z13-18:OH; Z13-18:Ald) and two other insects’ pheromone compounds (Z9-16:Ald; Z11-16Ald; Z11-16:OH;Z11-16:Ac and 16:Ald). The results indicate *C. medinalis* can detect those pheromones and is sensitive to Z11-16:Ald and Z9-16:Ald. In the heterologous expression system of Xenopus oocytes, we cloned three pheromone receptor genes, CmedPR1, CmedPR2, and CmedPR3. These had the same electroantennogram response, in addition to the response to the conspecific pheromone. CmedPR2 and CmedPR3 displayed strong sensitivity to Z11-16Ald and Z9-16:Ald. These results may contribute to clarifying how *C. medinalis* recognizes pheromones and interspecies communication.

## Introduction

Olfactory receptors profoundly influence insects. Many insect behaviors are affected by their olfactory receptors during various processes such as seeking mates, finding host plants and oviposition. Olfactory receptors, located in sensory neuron membranes, help insects sense compounds in their environment. After more specific study, we divided olfactory receptors into two types, odorant receptors (ORs) and pheromone receptors (PRs). ORs primarily perceive plant and floral scents, and PRs are responsible for sensing insect pheromones released by female insects (Raina 1989). The *C. medinalis* sex pheromone has four components, (Z)-11-octadecenal (Z11-18:Ald), (Z)-13-octadecenal (Z13-18:Ald), (Z)-11-octadecen-1-ol(Z11-18:OH), and (Z)-13-octadecen-1-ol (Z13-18:OH) (Kawazu et al. 2000; Rao 1995). *C. suppressalis* and *S. inferens* are two common insects jeopardizing rice crops. Their sex pheromone components were identified in previous studies (Nagayama et al. 2006; Nesbitt et al. 1975; Tatsuki et al. 1983; Wu and Cui 1986; Zhu et al. 1987). Notably, some studies showed that females detect sex pheromones to avoid places of high mating competition and unfavorable oviposition sites, thereby minimizing competition for ecological resources (Harari and Steinitz 2013; Holdcraft et al. 2016). Exposure of the female to the sex pheromone of its own species stimulates oviposition for some moth species, e.g., *Choristoneura fumiferana* (Palanaswamy and Seabrook 1978), but deters oviposition by other moth species (Goekce et al. 2007; Harari et al. 2011; Palanaswamy and Seabrook 1978). Females of two noctuids, *Heliothis armigera* (Hübner) and *Helicoverpa zea* (Boddie) were significantly repelled by their pheromones in olfactometer tests (Saad et al. 1981). In general, males compete for access to females whereas females compete for access to resources (Rubenstein 2012; Tobias et al. 2012).

Genome sequences related to *C. medinalis* were first published in 2012 (Shang et al. 2012). CmedORco (Liu et al. 2013a) and two Sensory neuron membrane proteins (SNMPs) genes (Liu, Su, et al. 2013b) were subsequently cloned. Zeng et al. (2015) systematically analyzed the chemical sense gene, including 24 ORs, 4 PRs, 15 Ionotropic receptor(IRs), 30 Odorant binding proteins, (OBPs), 26 chemosensory proteins (CSPs), and two SNMPs. Liu (2017) added 29ORs, 15 IRs, 12OBPs, 15 CSPs, and two SNMPs to the newly discovered olfactory genes. During experiment implementation, however, we found 4 CmedPRs should have been 3. PR4 is part of PR1 rather than a new PR. In addition, not all the CmedPRs submitted were completed sequences although Zeng thought the sequences were complete. We cloned a full sequence of three PRs based on their sequences. In the identity function of Cmed olfactory genes, most experiments were conducted in OBPs (Sun et al. 2019) and CSPs (XS, G, Duan et al. 2019). Olfactory genes related to pheromones are CmedPBP4, which are the Z13-18:Ac, Z11-16:Al, and Z13-18:OH binding proteins (Sun et al. 2016) and CmedCSP3, which has high binding affinities to Z11-16:Ac and Z11-16:Al (Zeng et al. 2018).

In recent decades, with the applications of transcriptome technology, many pheromone receptors have been deorphaned by vivo heterologous expression systems. These are (i) Xenopus oocytes coupled with voltage–clamp electrophysiology, (ii) mammalian or insect cell lines coupled with calcium imaging, and (iii) the so-called Drosophila “empty neuron” and T1 sensillum systems in combination with electrophysiological single sensillum recordings (Fleischer et al. 2018; Forstner et al. 2009; Grosse-Wilde et al. 2006; Kurtovic et al. 2007; Mitsuno et al. 2008; Nakagawa et al. 2005; Pask et al. 2017; Syed et al. 2010 reviewed in Montagne et al. 2015). For the assessment of candidate PRs (and other ORs), the Xenopus oocyte system has been most widely applied (Liu et al. 2017; Nakagawa et al. 2005; Sun et al. 2013; Zhang et al. 2015).

In previous studies, pheromone receptors were only identified as conspecific sex pheromones (Mitsuno et al. 2008; Sun et al. 2013; Zhang, 2010), but some studies found a part of PRs recognized that some compounds do not belong to their pheromone components (Chang et al. 2015; Liu et al. 2018; Zhang et al. 2014; Zhang et al. 2019; Zhang and Löfstedt 2013). In our study, we systematically tested three PR genes of *C. medinalis* using conspecific sex pheromones and other pheromone compounds belonging to two common insects living in rice fields, *C.suppressalis* and *S. inferens*. Test results indicated that *C. medinalis* detected the pheromones of three insects and have a strong sensitivity to them. The test revealed pheromone receptors not only have functions for seeking mates but also help insects perceive other species’ pheromones in the same ecological environment, which may help them avoid competition for food.

## Materials and Methods

### 1. Insect tissue collection

*C. medinalis* were collected from a rice field in XiaoShan Hangzhou, China (N30°18 ‘12.93” E120°34’ 24.75”). Antennae of *C. medinalis* were cut off on a sterile operating table and then immediately frozen in liquid nitrogen and stored at −70°C until use.

### 2. Electroantennogram recordings

The electrophysiological recordings of whole male and female antennae in response to conspecific sex pheromone components and other species pheromone compounds were conducted according to the standard technique (Cao et al. 2016). The components used in the EAG assay were dissolved in paraffin oil and diluted to 10 μg/μL. A piece of filter paper (0.5 × 5 cm) loaded with 10 μL pheromones was used as a stimulus, and paraffin oil was used as a control. Moths captured from the field were tested, and signals from antennae were amplified with a 10 × AC/DC headstage preamplifier (Syntech, Kirchzarten, Germany) and further acquired with an Intelligent Data Acquisition Controller (IDAC-4-USB; Syntech, Kirchzarten, Germany). Signals were recorded using SyntechEAG-software (EAGPro 2.0).

### 3. RNA isolation and RACE amplification

Total RNA was extracted from collected tissues using TRIzol reagent (Invitrogen, Ambion, USA). The RNA extraction method followed the manufacturer’s instructions. The single-stranded cDNA templates were synthesized with 1 μg antennal total RNA using SuperScript™ III First-Strand Synthesis System (Thermo Fisher Scientific, USA). Based on the partial sequence previously identified by analyzing the transcriptome data of *C. medinalis* antennae (Zeng et al. 2015), primers were designed to amplify the core sequence by RT-PCR. Using the core sequences obtained from sequencing results and a 5’/3’ RACE KIT, Second Generation (Roche), we obtained the full-length sequence of the 3 CmedPR genes.

### 4. Quantitative PCR

The relative expression levels of CmedPRs and CmedORco transcripts in male and female antennae were compared using qPCR on a CFX connect real-time system (Bio-Rad USA). The qPCR reactions (25 μL) contained 12.5 μL of TB Green® Premix Ex Taq™ II (Takara, Japan), 0.5 μL of each primer (10 μM), 1 μL of cDNA, and 9 ul Easy Dilution (Takara, Japan). The cycling parameters were an initial denaturation at 95°C for 30 s, followed by 39 cycles of 95°C for 5 s and 60°C for 30 s. The actin gene and tubulin were used to verify the integrity of the cDNA templates (GenBank JN029806.1) (Li et al. 2012). The experiment was repeated 3 times using 3 independent RNA samples. Gene expression levels were analyzed using the 2-ΔΔCT method, where ΔCT = CT PR gene – CT actin gene and Δ Δ CT = ΔCT different tissues – ΔCT maximum (Livak and Schmittgen 2001).

### 5. Vector construction and cRNA synthesis

The specific primers with the Kozak consensus sequence and pT7Ts homologous sequence restricted by ECORV enzyme were designed to amplify the full open reading frames of the three CmedPRs and then be infusion into an expression vector pT7Ts The extracted plasmids were linearized by digestion with EcoRI and used as templates to synthesize cRNAs using the mMESSAGE mMACHINE T7 Kit (Ambion, USA). The purified cRNAs were diluted with nuclease-free water at a concentration of 2 μg/μL and stored at −80°C.

### 6. Electrophysiology and data analysis

We selected mature, healthy oocytes (Stages V–VII) and then added them to Wash Buffer (96mM NaCl, 2mM KCl, 5Mm MgCl2, and 5Mm HEPES) with Liberase TM Research Grade (1.5mg/ml) (Roche, USA) for 15 min at room temperature. We microinjected 27.6ng CmedORco and CmedPR genes into the oocytes. After injection, the oocytes were cultured for 4–7 days at 1°C in 1× Ringer’s solution (96 mM NaCl, 2 mM KCl, 5 mM MgCl2, 0.8 mM CaCl2, and 5 mM HEPES [pH 7.5]) supplemented with 5% dialyzed horse serum, 50 mg/ml tetracycline, 100 mg/ml streptomycin, and 550 mg/ml sodium pyruvate. We recorded whole-cell currents from the injected Xenopus oocytes with a two-electrode voltage clamp and used an OC-725C oocyte clamp (Warner Instruments, Hamden, CT, USA) at a holding potential of −80 mV. Voltage clamp data recording and analysis were completed using Digidata 1550B and Pclamp10.0. Curves were analyzed using GraphPad Prism 8.0 (GraphPad Software Inc., San Diego, CA, USA).

## Results

In this study, 4 sex pheromone components and 4 pheromone analogs were chosen to evaluate the antennal EAG responses of male and female *C. medinalis*. The results showed that all tested compounds elicited EAG responses of male antennae at a dose of 100 μg. Z11-16:Ald evoked the strongest EAG responses from the antennae of male moths. The EAG responses of conspecific pheromone components Z13-18:Ald, Z11-18:Ald and interspecific pheromone compounds Z11-16:Ac, Z9-16:Ald all exceeded 2 mV (Fig. 1).

**Figure 1.**
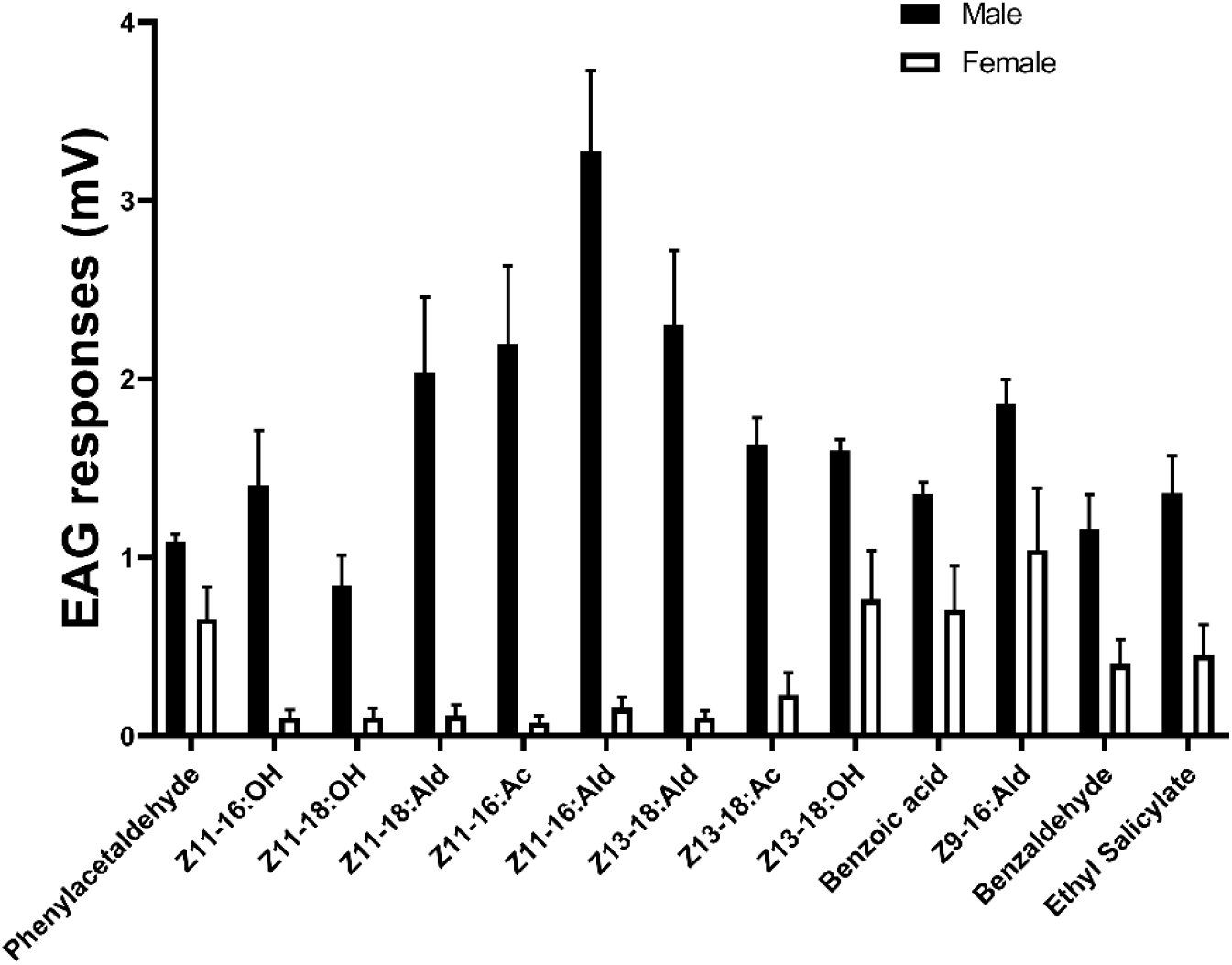
Electroantennography (EAG) responses from antennae of male and female C. medinalis to pheromone compounds and odorants.

We used qPCR to compare three PR genes’ expression in different tissues and genders. Three PR genes were expressed primarily in male *C. medinalis*. Interestingly, three PRs have different expression levels in female insects but are nearly undetectable in other tissues, except for antennae (Fig. 2).

**Figure 2.**
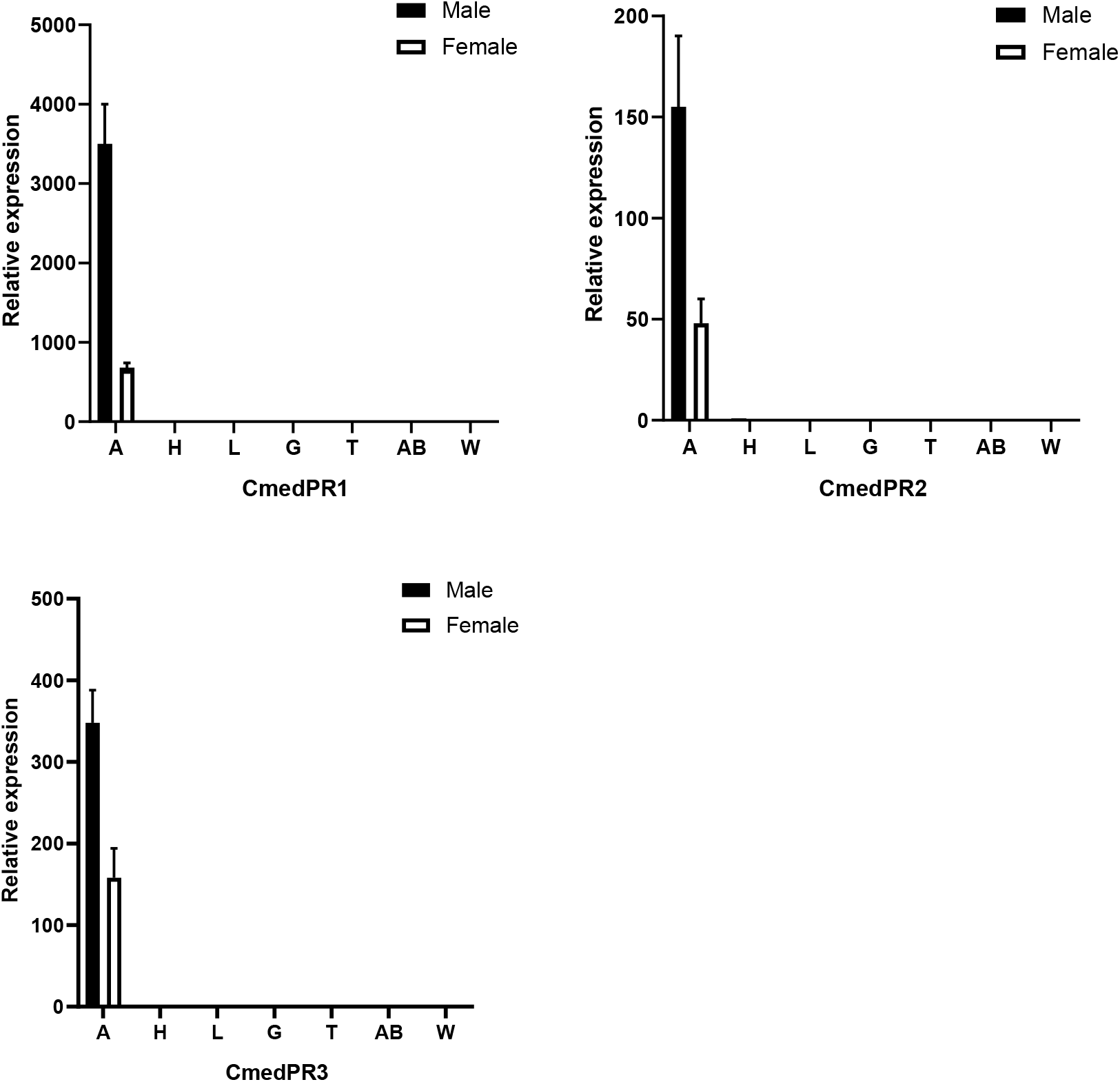
Tissue- and sex-specific expression of the *C. medinalis*. Expression of *C. medinalis*. PRs in 7 tissues of 2 sexes, including antennae (A), heads (H), legs (L), genitals (G), thoraxes (T), abdomens (AB), and wings (W). Error bars indicate SE.

The average values of expression levels of all sex pheromone receptor genes are from three biological replications by quantitative real-time PCR. All sex pheromone receptor genes were expressed in male antennae with higher levels than in female antennae. Among the 3 genes, CmedPR1 showed the highest expression level in male antennae, ~3.5- and 4-fold higher than in female antennae (Fig. 2).

### Responses of PRs to pheromone components of *C. medinalis*

The functional characterization of the 3 PRs was accomplished using Xenopus oocyte expression and the voltage clamp recording system. Each of the three pheromone receptors were co-expressed with the CmedORco in Xenopus oocytes for 4–6 days. Testing pheromone components included Z11-18:OH, Z11-18:Ald, Z13-18:Ald, and Z13-18:OH. CmedPR1 responded to octadecenol and octadecenal that had 11C unsaturated (15 ± 1.5nA and 45 ± 2nA, respectively). CmedPR2 responded to octadecenol and octadecenal that had 13C unsaturated (46 ± 1.6nA and 106 ± 6.4nA, respectively). CmedPR3 co-identified two octadecenol components with CmedPR1 and CmedPR2. Response currents were up to 84 ± 5.9nA (Z13-18:OH) and 165.5 ± 9.5nA (Z11-18:OH) (Fig. 3).

**Figure 3.**
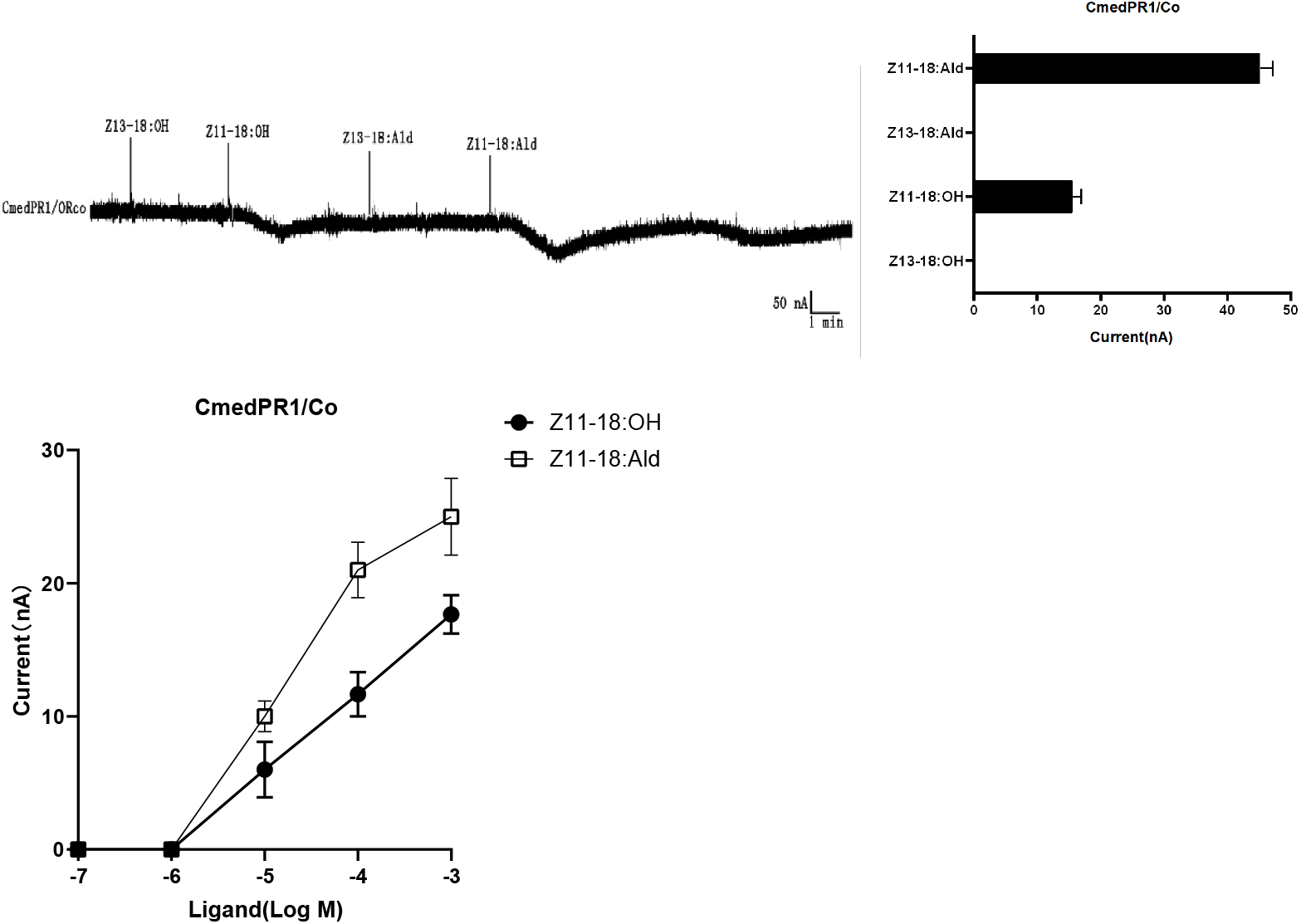
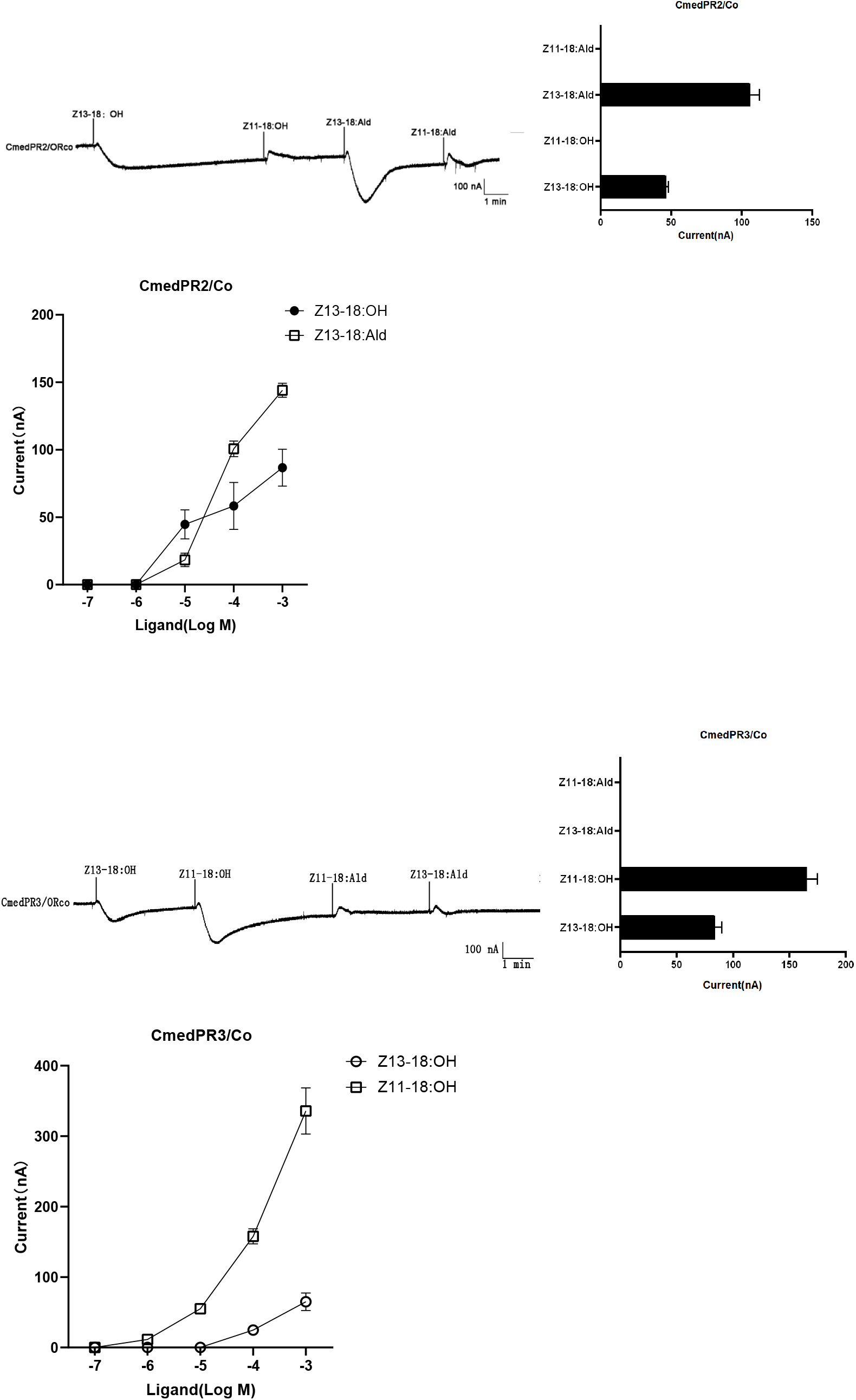
Responses of Xenopus oocytes with co-expressed CmedPRs/ORco to stimulation with pheromone compounds. In each panel: (Left) Inward current responses of CmedPR/ORco co-expressed Xenopus oocytes to 10-4 mol/L sex pheromone components and analogs. (Right) Response spectrum of PRs. (Bottom) Responses of CmedPRs at different doses of each stimulus. Error bars indicate SEM (n = 3).

### Responses of PRs to pheromone components of other Lepidopteran insects and analogues

To further confirm communication between *C. medinalis* and other Lepidopteran insects in rice fields, we tested several kinds of pheromone components and analogues. The results indicated that 3 CmedPRs respond to some pheromone components, all of which belong to *Chilo suppressali* and *Sesamia inferens*.

CmedPR1 had a weak response to Z11-16:Ac and 16:Ald. Interestingly, CmedPR2 and CmedPR3 both identified function in Z11-16:Ald and Z9-16:Ald. In addition, the response current to these components exceeded 140 nA, even reaching up to 250 nA (CmedPR3,Z9-16:Ald), which is higher than in the self-pheromone components. Moreover, in dose-response studies, 10^-7^ M Z9-16:Ald and Z11-16:Ald could elicit significant responses from oocytes that co-expressed CmedPR2 and CmedPR3 with CmedORco. This shows that *C. medinalis* has greater sensitivity to these components than conspecific pheromone components (Fig. 4).

**Figure 4.**
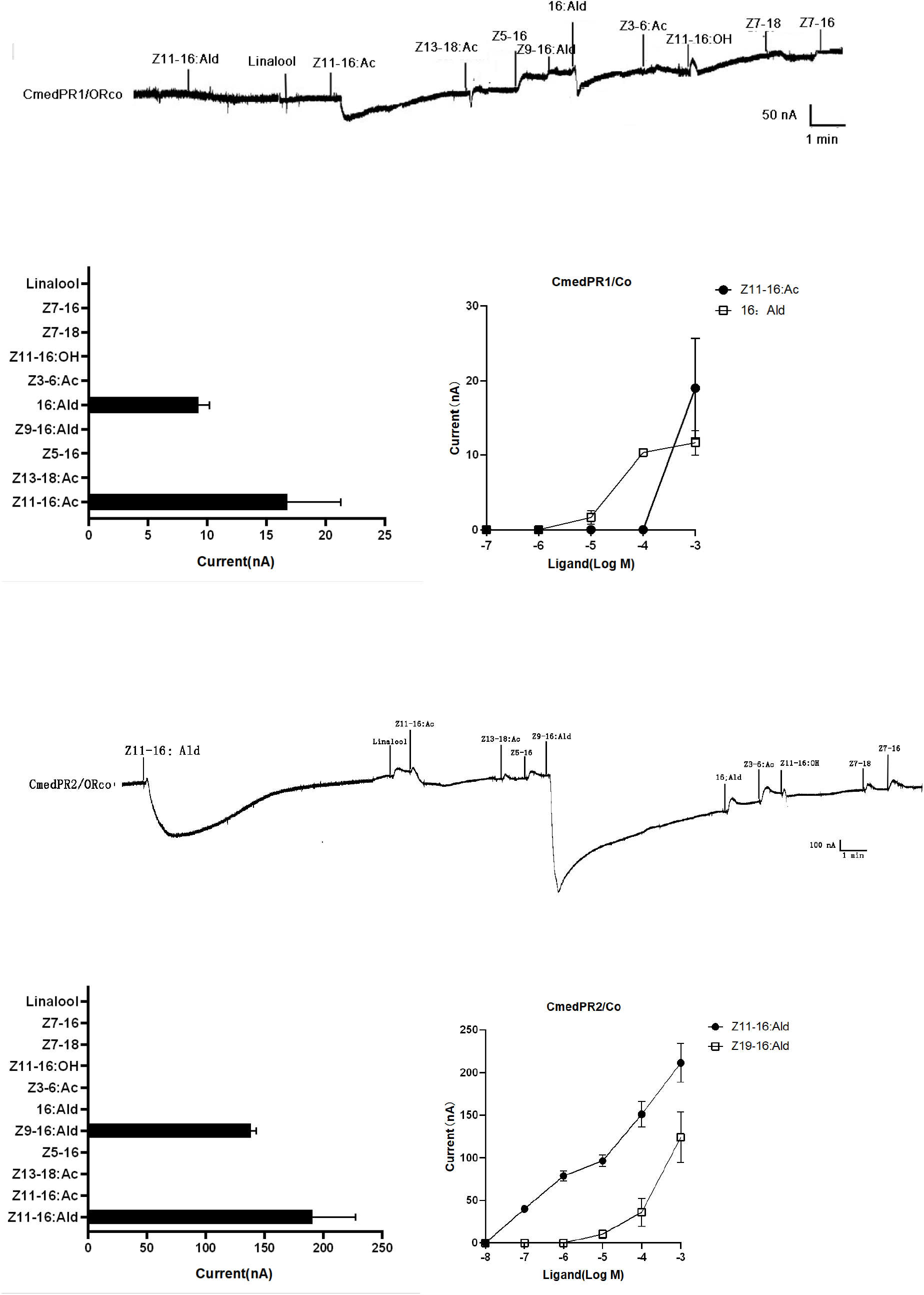
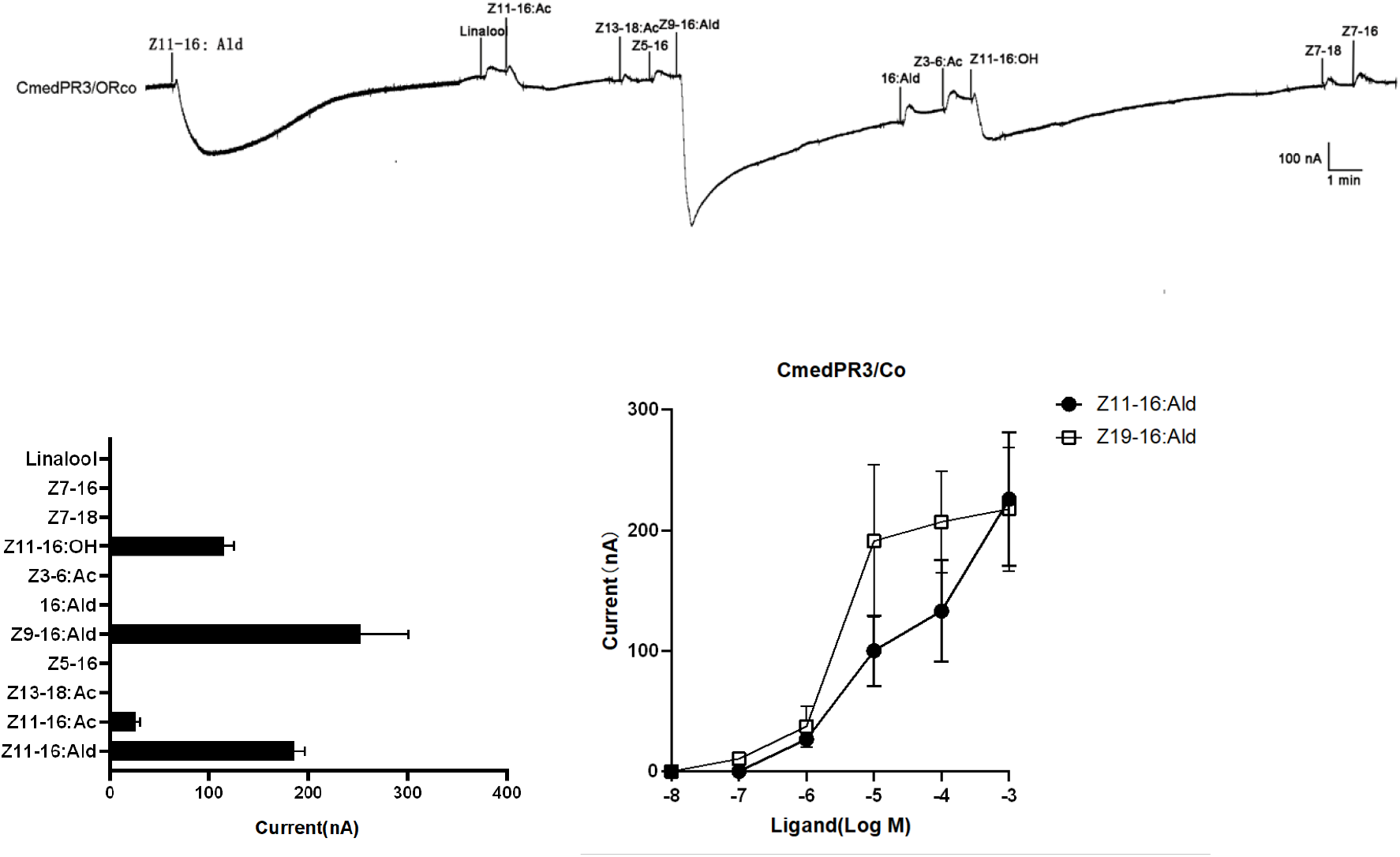
Responses of Xenopus oocytes with co-expressed CmedPRs/ORco to stimulation with pheromone compounds of other Lepidopteran insects and analogues. In each panel: (Top) Inward current responses of CmedPR/ORco co-expressed Xenopus oocytes to 10-4 mol/L sex pheromone components and analogs. (Left) Response spectrum of PRs. (Right) Responses of CmedPRs at different doses of each stimulus. Error bars indicate SEM (n = 3).

## Discussion

Although it is common for insects to sense other species’ pheromones and exhibit corresponding behavior, experiments identifying the function of pheromone receptors to verify those behaviors are rarely conducted. Previous studies in the olfactory system of *C. medinalis* reported indirect evidence of interspecific communication (Sun et al. 2016; Zeng et al. 2018).

In our study, we tested the EAG response of *C. medinalis* to conspecific pheromones and the pheromones of other species living in the rice field. The results indicate *C. medinalis* has sensitive responses to Z11-16:Ald and Z9-16:Ald, which do not belong to conspecific pheromones. Future studies may prove that *C. medinalis* can sense those pheromone compounds. In the meantime, 3 CmedPRs co-expressing with CmedORco in Xenopus oocytes further confirm that *C. medinalis* has strong responses to the pheromone compounds of *C.suppressalis* and *S. inferens*. The reason *C. medinalis* is so sensitive to these 2 insects may be the feeding habits of the 3 larvae. The larvae of *C. suppressalis* and *S. inferens* jeopardize the rice stalks and make the rice wither. This change strongly affects *C. medinalis* larvae’s feeding on rice leaves. Thus, *C. medinalis* may detect the existence of *C. suppressalis* and *S. inferens* to avoid food resource competition. This indicates that pheromone receptors have more important roles in interspecies communication than helping insects find mates. This study may also inspire other studies to determine whether pheromone receptors have ligands that do not belong to conspecific pheromones.

## References

Chang H, Liu Y, Yang T, et al. Pheromone binding proteins enhance the sensitivity of olfactory receptors to sex pheromones in Chilo suppressalis[J]. Scientific reports, 2015, 5: 13093.

females of both species. Entomol. Exp. Appl. 1981, 30, 123–127.

Forstner, M., Breer, H., and Krieger, J. (2009). A receptor and binding protein interplay in the detection of a distinct pheromone component in the silkmoth Antheraea polyphemus. Int. J. Biol. Sci. 5, 745–757. doi: 10.7150/ijbs.5.745

Goubault M, Cortesero AM, Poinsot D, Wajnberg E, Boivin G, 2007. Does host value influence female aggressiveness, contest outcome and fitness gain in parasitoids? Ethology 113(4):334–343.

Grosse-Wilde, E., Svatos, A., and Krieger, J. (2006). A pheromone-binding protein mediates the bombykol-induced activation of a pheromone receptor in vitro. Chem. Senses 31, 547–555. doi: 10.1093/chemse/bjj059

Harari AR, Zahavi T, Thiery D, 2011. Fitness cost of pheromone production in signaling female moths. Evolution 65(6): 1572–1582.

Harari, A. R., and Steinitz, H. (2013). The evolution of female sex pheromones. Curr. Zool. 59, 569–578. doi: 10.1093/czoolo/59.4.569

Holdcraft, R., Rodriguez-Saona, C., and Stelinski, L. L. (2016). Pheromone autodetection: evidence and implications. Insects 7:E17. doi: 10.3390/insects7020017.

Kurtovic, A., Widmer, A., and Dickson, B. J. (2007). A single class of olfactory neurons mediates behavioural responses to a Drosophila sex pheromone. Nature 446, 542–546. doi: 10.1038/nature05672

Li, Shang-Wei, et al. “Transcriptome and gene expression analysis of the rice leaf folder, Cnaphalocrosis medinalis.” PLoS One 7.11 (2012): e47401.

Liu W, Jiang X, Cao S, et al. Functional studies of sex pheromone receptors in asian corn borer Ostrinia furnacalis[J]. Frontiers in physiology, 2018, 9: 591.

Liu, F., Xiong, C., and Liu, N. (2017). Chemoreception to aggregation pheromones in the common bed bug, Cimex lectularius. Insect Biochem. Mol. Biol. 82, 62–73. doi: 10.1016/j.ibmb.2017.01.012

Liu, Su, et al. “Identification and characterization of two sensory neuron membrane proteins from Cnaphalocrocis medinalis (Lepidoptera: Pyralidae).” Archives of insect biochemistry and physiology 82.1 (2013): 29–42.

Liu, Su, et al. “Transcriptome sequencing reveals abundant olfactory genes in the antennae of the rice leaffolder, Cnaphalocrocis medinalis (Lepidoptera: Pyralidae).” Entomological Science 20.1 (2017): 177–188.

Mitsuno, H., Sakurai, T., Murai, M., Yasuda, T., Kugimiya, S., Ozawa, R., et al. (2008). Identification of receptors of main sex-pheromone components of three Lepidopteran species. Eur. J. Neurosci. 28, 893–902. doi: 10.1111/j.1460-9568.2008.06429.x

Nagayama, A., Wakamura, S., Taniai, N. and Arakaki, N. (2006) Reinvestigation of sex pheromone components and attractiveness of synthetic sex pheromone of the pink borer, Sesamia inferens Walker (Lepidoptera: Noctuidae) in Okinawa. Appl Entomol Zool 41: 399–404.

Nakagawa, T., Sakurai, T., Nishioka, T., and Touhara, K. (2005). Insect sexpheromone signals mediated by specific combinations of olfactory receptors. Science 307, 1638–1642. doi: 10.1126/science.1106267

Nesbitt, B. F., P. Beevor, D. Hall, R. Lester, and V. Dyck. 1975. Identification of the female sex pheromones of the moth, Chilo suppressalis. J. Insect Physiol. 21: 1883–1886.

Palanaswamy P, Seabrook WD, 1978. Behavioral responses of the female eastern spruce budworm Choristoneura fumiferana(Lepidoptera, Tortricidae) to the sex pheromone of her own species. J. Chem. Ecol. 4(6): 649–655.

Pask, G. M., Slone, J. D., Millar, J. G., Das, P., Moreira, J. A., Zhou, X. F., et al. (2017). Specialized odorant receptors in social insects that detect cuticular hydrocarbon cues and candidate pheromones. Nat. Commun. 8:297. doi: 10.1038/s41467-017-00099-1

Raina, A. K., H. Jaffe, T. G. Kempe, P. Keim, R. W. Blacher, H. M. Fales, C. T. Riley, J. A. Klun, R. L. Ridgway, and D. K. Hayes. 1989. Identification of a neuropeptide hormone that regulates sex pheromone production in female moths. Science. 244: 796–798.

Rao, A. Ganeswara, et al. “Identification and field optimisation of the female sex pheromone of the rice leaffolder, Cnaphalocrocis medinalis in India.” Entomologia experimentalis et applicata 74.3 (1995): 195–200.

